# Vitamin A supplement after neonatal *Streptococcus pneumoniae* pneumonia alters CD4^+^T cell subset and inhibits allergic asthma in mice model

**DOI:** 10.1101/412940

**Authors:** Yonglu Tian, Qinqin Tian, Yi Wu, Xin Peng, Yunxiu Chen, Qinyuan Li, Guangli Zhang, Xiaoyin Tian, Luo Ren, Zhengxiu Luo

**Affiliations:** Key Laboratory of Pediatrics in Chongqing; Department of Children’s Hospital of Chongqing Medical University of Education, Key Laboratory of Child Development and Disorders; Department of Respiratory Medicine, Children’s Hospital of Chongqing Medical University, Chongqing, China

**Keywords:** Neonatal, *S.pneumoiae* pneumonia, asthma, vitamin A

## Abstract

**Background:** Previously, we showed that neonatal pneumonia caused by *Streptococcus pneumoniae* (*S. pneumoniae*) promoted adulthood ovalbumin (OVA) induced allergic asthma. Many studies have demonstrated that vitamin A deficiency induced the development of allergic asthma. Whether neonatal *S. pneumoniae* pneumonia promoted allergic asthma development was associated with vitamin A concentrations remains unclear.

**Methods:** Female BALB/c neonates were infected with *S. pneumoniae* strain D39 and subsequently treated with vitamin A. Vitamin A concentrations in lung, serum and liver were monitored on 2, 5, 7, 14, 21, 28 days post infection. Four weeks after infection, mice were sensitized and challenged with OVA to induce allergic airway disease (AAD) in early adulthood. Twenty-four hours after the final challenge, lung histo-pathology, cytokine concentrations in bronchoalveolar lavage fluid (BALF), airway hyperresponsiveness (AHR) and lung CD4^+^T cells were measured.

**Results:** We demonstrated that neonatal *S. pneumoniae* pneumonia induce lung vitamin A deficiency up to early adulthood. Moreover, neonatal *S. pneumoniae* pneumonia aggravated airway inflammatory cells accumulation and increased AHR during AAD, decreased Foxp3^+^Treg and Th1 productions remarkably, while Th2 cell expression was increased significantly. Further study indicated that vitamin A supplement after neonatal *S. pneumoniae* pneumonia can promote Foxp3^+^Treg and Th1 productions, decrease Th2 cell expressions, alleviate AHR and inflammatory cells infiltration during AAD.

**Conclusions:** Using a mouse model, we demonstrate that Vitamin A supplement after neonatal Streptococcus pneumoniae pneumonia alters the CD4^+^T cell subset and inhibits the development of early adulthood allergic asthma.

## Background

Asthma is a heterogeneous disease, characterized by airway chronic inflammation together with airway hyperresponsiveness^[1]^. It is more common in childhood, and most adult asthma originate from childhood indicating that childhood events have an important role in asthma pathogenesis^[2–4]^. Childhood is an important period for the maturation of the immune system, specific infections may alter immunologic programming, which plays critical role in the progression of allergic airways disease (AAD)^[5]^. Neonatal infections caused by *Streptococcus pneumoniae* (*S. pneumoniae*), Haemophilus influenzae, Moraxella catarrhalis can increase the risk of bronchiolitis^[6]^ and preschool asthma^[7]^. *S. pneumoniae* is the most common bacterial pathogen of community acquired pneumonia in childhood. Our previous study suggested that neonatal *S. pneumoniae* pneumonia promoted OVA-induced asthma development^[8]^. Although the prevention and treatment of asthma induced by *S. pneumoniae* pneumonia is crucial, while it remains indistinctly. Pneumonia continues to be a serious health issue worldwide; affecting millions annually, increasing morbidity and mortality globally^[9–12]^. Pneumonia decreases vitamin A levels significantly in children under five years old^[13]^. Infections may affect vitamin A intake, absorption, storage, release, distribution and metabolism^[14]^. Vitamin A is predominantly stored in the liver as retinyl esters with lung being a secondary storage site. Evidence shows that vitamin A deficiency may be associated with asthmatic development^[15, 16]^. Our previous study indicated that the severity of vitamin A deficiency was associated with the course and severity of wheezing in infants^[17]^. Whether neonatal *S. pneumoniae* pneumonia induced adulthood allergic asthma was associated with vitamin A deficiency remains unclear. In this study, we established a neonatal non-lethal *S. pneumoniae* pneumonia mouse model and monitored vitamin A levels in lung, serum and liver until early adulthood. We explored the effects of vitamin A supplement after neonatal *S. pneumoniae* pneumonia on the development of adulthood allergic asthma. Our data demonstrated that neonatal *S. pneumoniae* pneumonia induced vitamin A deficiency in the lung up to early adulthood and vitamin A supplement altered the CD4^+^T cell subset and inhibited early adulthood allergic asthma development in mice.

## Methods

### Mice

Parturient BALB/C mice were purchased from Animal Resources Centre, Chongqing medical university. Pregnant mice were kept separately and monitored for births. Newborn female mice were raised in a pathogen-free environment, and housed at 24 °C;under a 12h light, 12h dark cycle, and given a normal diet and water. All experiments performed in mice were permitted by the Institutional Animal Care and Research Advisory Committee at the Chongqing Medical University. All experimental animals were used in accordance with the guidelines issued by the Chinese Council on Animal Care.

### Establishment of a Neonatal non-lethal *S. pneumoniae* pneumonia mouse model

Neonatal *S. pneumoniae* pneumonia (*S.pp*) was established according to the procedures described in our previous study. Briefly, *S. pneumoniae* (D39) was plated onto trypic soy broth (Pangtong, China), grown for 10-14 hours at 37°C in a 5% CO2 atmosphere, washed, and suspended in sterile phosphate buffered saline (PBS). Conscious neonatal (1-week-old) BALB/c mice were infected intranasally with 2×10^7^ CFU of *S. pneumonia* in 5ul of PBS. Mock-infected mice were injected intranasally with 5ul of PBS.

### Determination of Vitamin A concentrations in tissues

Lung, serum and liver were collected from uninfected controls and neonatal *S. pneumoniae* pneumonia mice on 2, 5, 7, 14, 21, 28 days post infection. After grinding, the liver and lung were extracted in ethane. Thereafter the extracted samples and the untreated serum were degassed and redissolved. Total retinol concentration in lung, liver and serum was determined by high performance liquid chromatograph (HPLC, Model G1315 A, Agilent Technologies, Palo Alto) using trimethylmethoxyphenyl-retinol as an internal standard^[18]^.

### Establishment of a Vitamin A supplement model

Retinyl palmitate (Sigma) with all-trans retinoic acid (Sigma) in the ratio of 10:1 was dissolved in rapeseed oil to configure to the vitamin A used for subsequent experiments^[19]^. 24 hours after post infection, neonates were administrated orally with a dose of 20^IU^/g of vitamin A once daily for four consecutive days^[20]^ to build the vitamin A supplement model.

### Induction of Allergic airway disease (AAD)

Four weeks after neonatal *S. pneumoniae* pneumonia infection (mice have matured into early adulthood in four weeks), mice were divided into the following groups: uninfected non-allergic (control), uninfected allergic (OVA), infected allergic (*S.pp* /OVA), infected vitamin A supplement allergic (*S.pp*+VA/OVA). To induce AAD, mice in the OVA, *S.pp* /OVA and *S.pp*+VA/OVA groups were sensitized with i.p. injections of 100 μg OVA (Sigma-Aldrich, St. Louis, MO, USA) diluted in 50% aluminum hydroxide gel (Sigma-Aldrich) for a total volume of 200 μL on days 35 and 42. From days 49-52, mice were exposed to 1% OVA aerosols for 30 min/d. Controls were simultaneously sensitized and challenged with sterile PBS. AAD was assessed within 24 h after the final challenge (Fig 1). Each experiment was repeated three times with a sample size of a total of four to eight mice per group.

**Fig 1.**
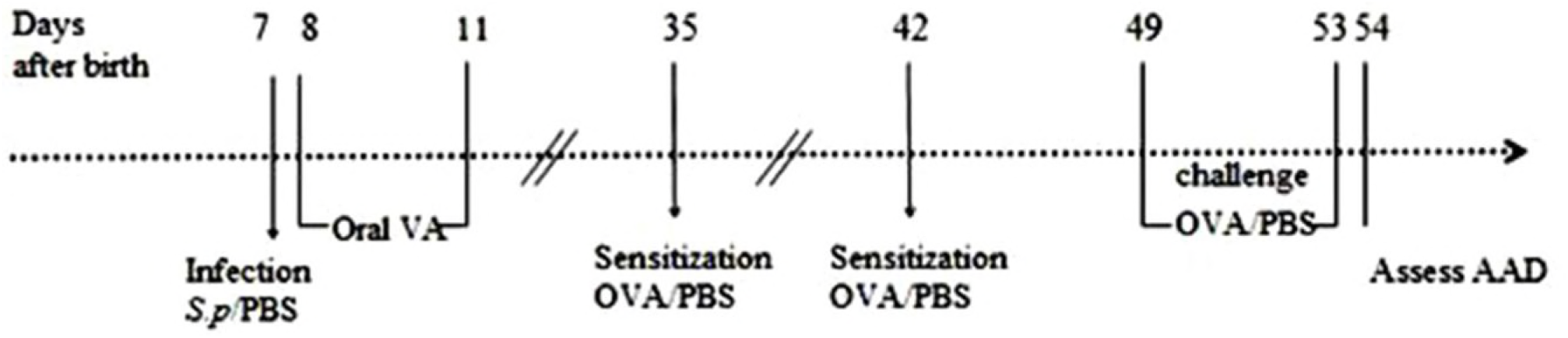
Establishment of models and schematic of study protocol. Neonatal *S. pneumoniae* pneumonia BALB/c mice were divided into the following groups: uninfected, non-allergic(Control), uninfected, allergic(OVA), neonatal infected, allergic(*S.pp /*OVA) and vitamin A supplement after neonatal infected, allergic (*S.pp*+VA/OVA). Mice were infected intranasally with *S. pneumoniae* or phosphate-buffered saline (PBS) on day 7 (1 week-old), and supplemented orally with vitamin A on days 8-11. Mice were sensitized by an i.p. injection of ovalbumin (OVA) or PBS on days 35 and42, and challenged with aerosolized OVA or PBS to induce allergic airways disease (AAD) from 49 to 52 days.

### Measurement of airway hyperresponsiveness (AHR)

AHR was assessed in vivo by measuring the changes in transpulmonary resistance using a mouse plethysmograph and methods previously described^[21–23]^. Briefly, 24 hours after the final challenge, AHR was measured in conscious, unrestrained mice by whole-body plethysmography (Emca instrument; Allmedicus, France). Each mouse was exposed to aerosolized PBS followed by increasing concentrations of aerosolized methacholine (Sigma-Aldrich, St. Louis, Mo. USA) solution (3.125, 6.25, 12.5, 25, and 50 mg/ml; Sigma) in PBS for 3 min and then rested for 2 min. The average Penh for each concentration was calculated from the continuously recorded pressure and flow data for 5 min. Penh is a dimensionless value and correlates with pulmonary airflow resistance. It represents a function of the ratio of peak expiratory flow to peak inspiratory flow and a function of the timing of expiration.

### Bronchoalveolar lavage fluid and cell counting

Twenty-four hours after the final challenge, mice were anesthetized with 10% chloral hydrate (0.1 mL/100 g, i.p). Bronchoalveolar lavage fluid (BALF) was obtained by flushing the lungs twice with 1 ml each of PBS through a cannulated trachea. The two aliquots were then pooled to obtain one sample for each mouse. Erythrocytes were lysed, and the remaining cells were centrifuged at 3000 rpm for 5 min. Total cell numbers in the BALF were determined using a standard hemocytometer. Differential cell counts were performed based on standard morphological and staining characteristics of at least 250 cells per sample. Supernatants were stored at –80 °C. All slides were characterized by a single blinded examiner to eliminate bias.

### Histo-pathology of lungs

Twenty-four hours after the final challenge, mice were euthanized by an intraperitoneal injection of a lethal dose of 10% chloral hydrate (0.3 mL/100 g, i.p.) to harvest the lungs. After fixing in formaldehyde for 24 hours, lungs were dissected and embedded in paraffin. Four micron thick sections were stained with hematoxylin and eosin (H&E; Sigma-Aldrich). At least five bronchi were selected from each mouse based on size (150-350mm in diameter) for analysis. The degree of airway inflammatory cell infiltration was scored in a single-blind fashion to reduce evaluator bias. Lung lesions were scored semi-quantitatively using a measurement tool as previously described^[24]^. Images were captured under a Nikon Eclipse E200 microscope connected to a Nikon Coolpix 995 camera (Nikon, Tokyo, Japan). The severity of inflammation was evaluated by assigning a value of 0 point for normal; 1 point for few cells; 2 points for a ring of inflammatory cells 1 cell layer deep; 3 points for a ring of inflammatory cells 2 to 4 cells deep; 4 points for a ring of inflammatory cells of >4 cells deep.

### BALF cytokines measurements

Concentrations of IL-4, IL-5, IL-13, IL-17A interferon (IFN)-γ and TGF-β (Xin Bosheng, Shenzhen, China) in BALF were detected by commercially available enzyme-linked immunosorbent assay (ELISA) kits according to the manufacturer’s instructions.

### Flow cytometric analysis of lung CD4^+^T cells

Lungs were minced and incubated 1 mL of RPMI 1640 containing 0.2% collagenase I (Sigma-Aldrich) for 15 min at 37°C. Single cell suspension was obtained by forcing tissue through a 70 μm cell filter (Becton, Dickinson and Company, Franklin Lakes, NJ, USA) After centrifugation, 3ml erythrocyte lysis buffer was added to the sediment. Fifteen minutes later, the cells were then harvested and washed and divided into two aliquots. One aliquot was stained for surface-associated CD11c-FITC (Rat anti-mouse; EB Biosciences) and CD4-FITC (Rat anti-mouse, BD Biosciences), CD25-PE (Rat anti-mouse, BD Biosciences.), Foxp3-PEcy5 (Rat anti-mouse, BD Biosciences) and the other was resuspended in RPMI 1640 medium containing 10% fetal bovine serum. The resuspended cells were incubated for 4–6 h at 37°C and 5% CO2 in 15 ml centrifuge tube in 1 mL medium containing phorbol 12-myristate 13-acetate (50 ng/mL; Sigma-Aldrich), ionomycin (500 ng/mL; Sigma-Aldrich) and GolgiPlug-containing brefeldin A (Becton, Dickinson and Company). To detect the subsets of Th1 and Th2 cells in lungs, cells were stained for intracellular IFN-γ-PerCP-Cy5.5 (Rat anti-mouse; Pharmingen), IL-17A-PE (Rat anti-mouse; Pharmingen), IL-4-APC (Rat anti-mouse; Pharmingen). Stained cells were detected by flow cytometry (FACS Canto; Becton, Dickinson and Company) and data were analyzed with CellQuest software (Becton, Dickinson and Company).

### Statistical Analysis

Results were analyzed using GraphPad Prism (version 5.0; GraphPad, La Jolla, CA, USA) and values are expressed as mean ± standard error. Statistical analysis was performed by either one-way analysis of variance (ANOVA) with Tukey’s post-test or two-way ANOVA with Bonferroni’s post-test. A value of P < 0.05 was considered significant.

## Results

### Neonatal *S. pneumoniae* pneumonia significantly decreases lung vitamin A levels in BALB/c mouse model

To assess if neonatal *S. pneumoniae* pneumonia(*S.pp*)caused vitamin A deficiency Vitamin A levels were measured in lung, serum and liver post-infection by HPLC. Results showed that the pulmonary vitamin A levels were significantly decreased in *S.pp* group as compared with the control group till early adulthood (Fig 2A). Serum vitamin A levels showed significant decline seen within the initial 2 weeks normalized over the next 2 weeks after pneumonia (Fig 2B). Vitamin A levels in liver were similar between neonatal *S. pp* and control groups (Fig 2C). These findings show that neonatal *S. pp* causes lung vitamin A deficiency in murine lungs.

**Fig 2.**
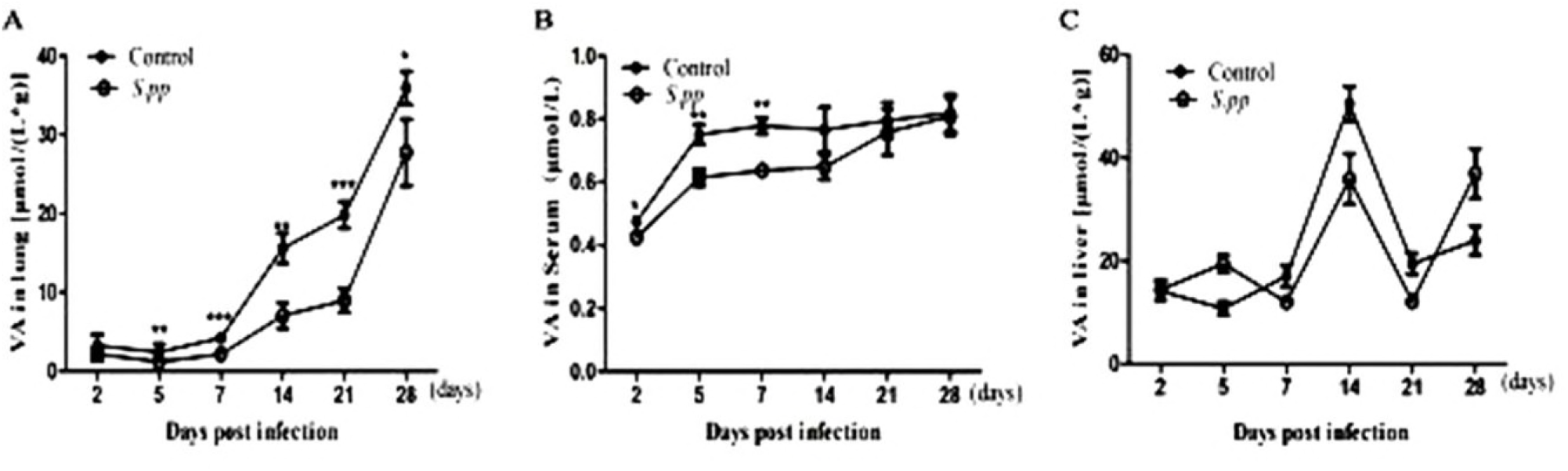
Neonatal *S.pp* effects on vitamin A status in lung (A), serum (B) and liver (C) on 2,5,7, 14, 21, 28 days post infection in BALB/C mice.*P<0.05, **P<0.01, ***P<0.001 as compared with the mock-infected (control) group, n=7 mice/group.

### Vitamin A supplement after neonatal *S. pneumoniae* pneumonia suppressed inflammatory cells infiltrate during AAD

Twenty-four hours after the final challenge, the total inflammatory cells, eosinophils and lymphocyte in the BALF from the OVA group were higher than that in the control mice. Interestingly, accumulation of total inflammatory cells, neutrophils, eosinophils, macrophages and lymphocyte in *S.pp*/OVA group were significantly increased as compared with the OVA group. In contrast, the number of total inflammatory cells, neutrophils, eosinophils, macrophages and lymphocyte was dramatically reduced in *S.pp*+VA/OVA group as compared with *S.pp*/OVA mice (Fig 3A-E). Our results clearly demonstrate that vitamin A supplement after neonatal *S. pp* significantly reduced inflammatory cells infiltration during AAD.

**Fig 3.**
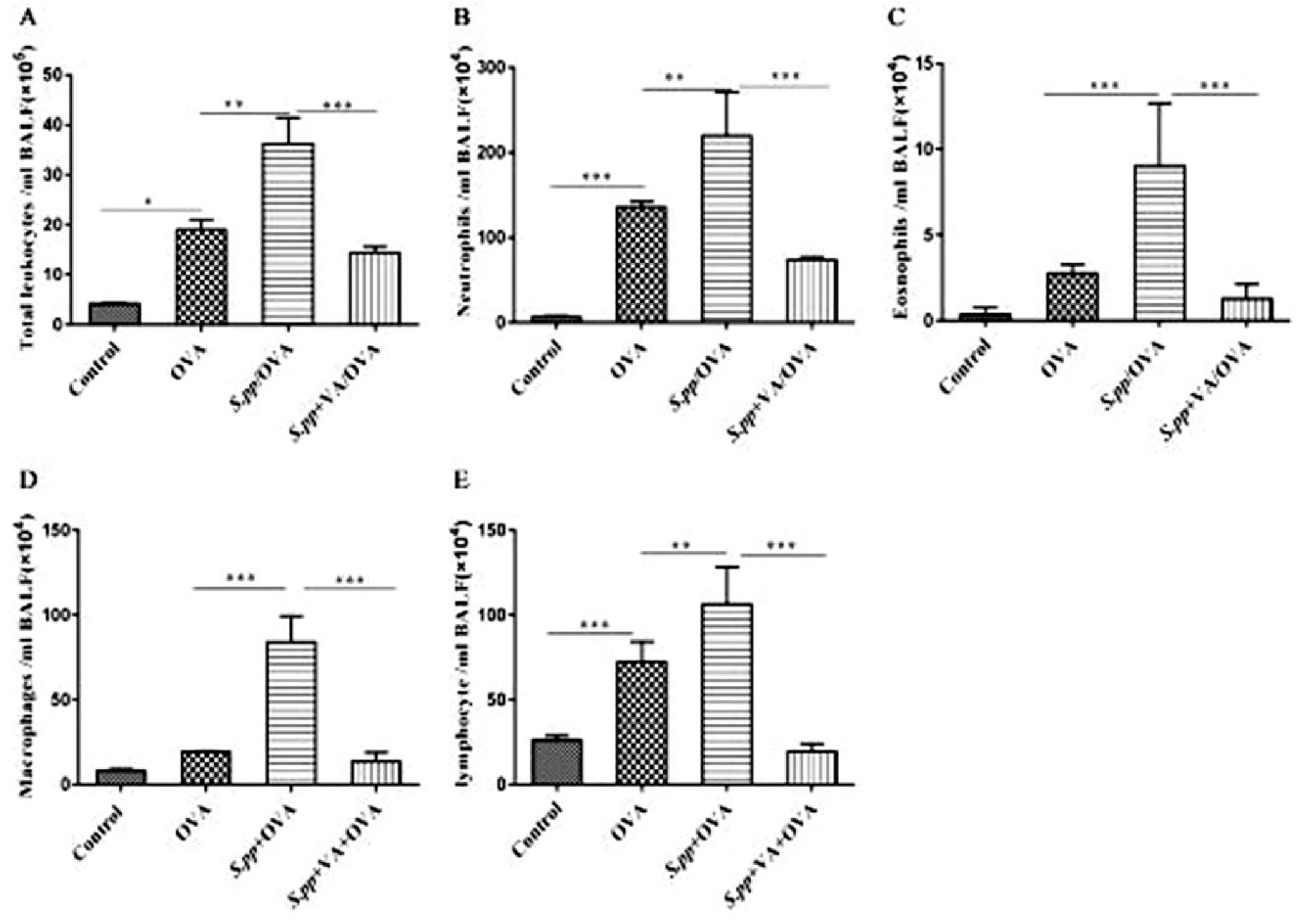
Vitamin A supplement after neonatal *S. pneumoniae* pneumonia significantly reduced inflammatory cells infiltration during AAD. Total cells (A), neutrophils (B), eosinophils (C), macrophages (D) and lymphocyte(E) were counted from bronchoalveolar lavage fluid (BALF) collected 24h after the final challenge. Control (uninfected, non-allergic); OVA (uninfected, allergic); *S.pp*/OVA (neonatal infected, allergic); *S.pp*+VA/OVA (vitamin A supplementary after neonatal infection, allergic). Data are shown as mean ± standard error from three separate experiments (n = 6–8 mice/group). *P<0.05, **P<0.01, ***P<0.001.

Histopathology of the lungs demonstrated neonatal *S. p*p significantly increased the accumulation of inflammatory cells around pulmonary alveoli, bronchioles and pulmonary vascular during AAD as compared with the control group. However, there were fewer inflammatory cells around pulmonary alveoli, bronchioles and pulmonary vascular in *S.pp*+VA/OVA group as compared with *S.pp*/OVA mice (Fig.4A-B). The inflammation scores of pulmonary peribronchitis, perivasculitis and alveolitis in the *S.pp* /OVA group were significantly higher than the OVA group. In contrast, the inflammation scores in *S.pp*+VA/OVA group were lower than the *S.pp* /OVA mice (P<0.05) (Fig 4 C-E). Taken together, these results clearly demonstrate that vitamin A supplement after neonatal *S. pp* significantly reduces infiltration of inflammatory cells during AAD.

**Fig 4.**
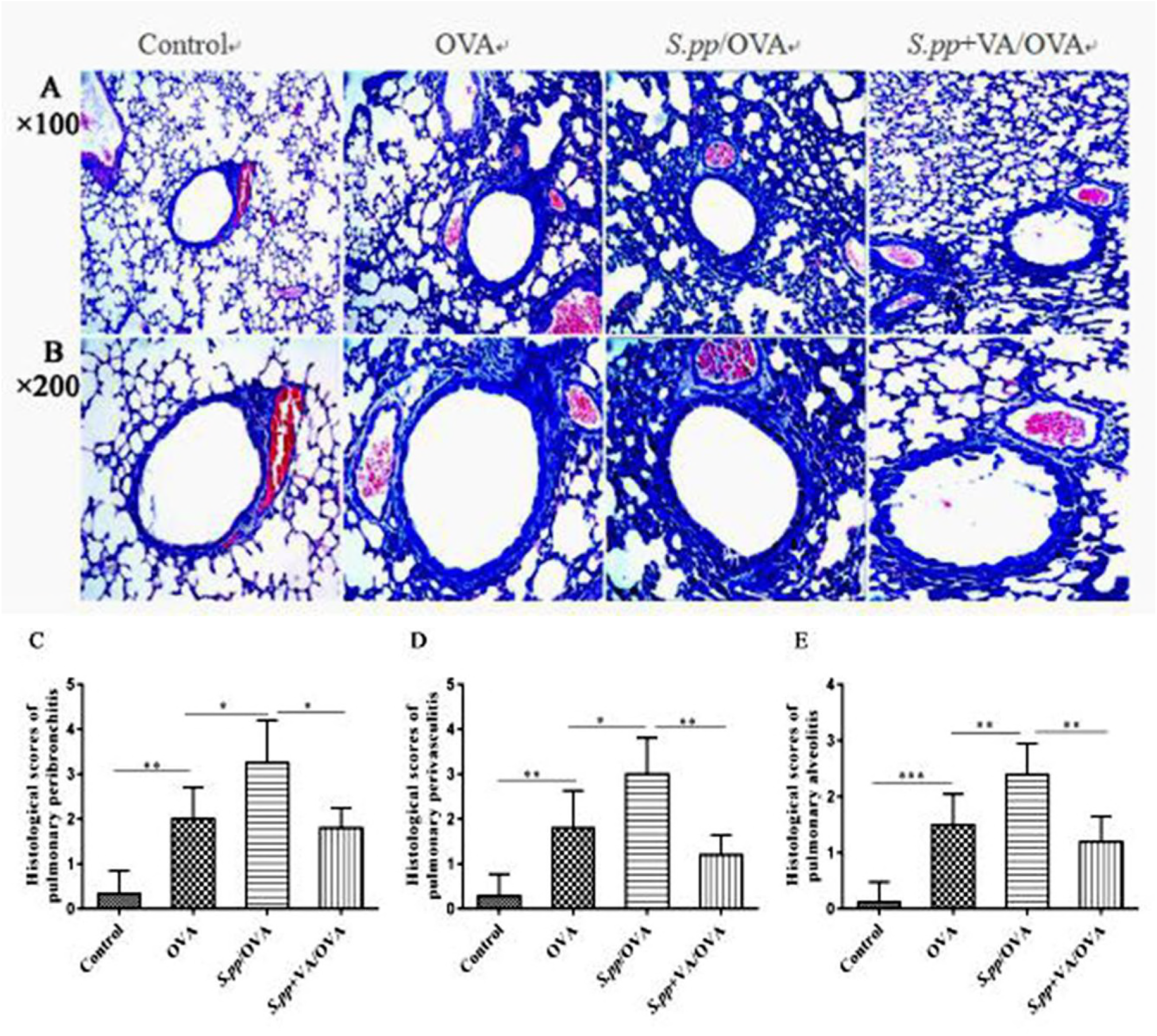
Vitamin A supplement after neonatal *S. pneumoniae* pneumonia significantly reduced lung inflammation during AAD. Hematoxylin and eosin (H&E) staining of lung tissue sections from uninfected, non-allergic (Control), uninfected, allergic (OVA), neonatal infected, allergic (*S.pp* /OVA) and vitamin A supplementary after neonatal infection, allergic (*S.pp* +VA/OVA) mice. Magnification: A×100, B×200. Histological scores of pulmonary peribronchitis (C), pulmonary perivasculitis (D) and pulmonary alveolitis (E). Data are shown as mean ± standard error from three separate experiments (n = 6–8 mice/group).*P<0.05, **P<0.01, ***P<0.001.

### Vitamin A supplement after neonatal *S. pneumoniae* pneumonia decreases AHR during AAD

Twenty-four hours after the final challenge, AHR was assessed by calculating the Penh values (i.e, enhanced respiratory pausing). Treatment with OVA remarkably increased AHR. The Penh value in *S.pp*+OVA group was significantly higher than the OVA group at methacholine concentrations of 12.5mg/ml (4.58±1.77*vs* 2.08± 0.59, P<0.001), 25mg/ml (4.76±1.43 *vs* 2.38± 0.55, P<0.001) and 50.0mg/ml (6.36±1.53 *vs* 2.74± 0.61, P<0.001). However, the Penh value in *S.pp*+VA/OVA group was significantly lower than *S.pp* /OVA group at methacholine concentrations of 12.5mg/ml (2.63±0.94 *vs* 4.58±1.77, P<0.001), 25mg/ml (3.06±0.94 *vs* 4.76±1.43, P<0.001) and 50.0mg/ml (2.90±1.01 *vs* 6.36±1.53, P<0.001).Vitamin A supplement after neonatal *S. p*p remarkably decreased AHR during AAD (Fig 5).

**Fig 5.**
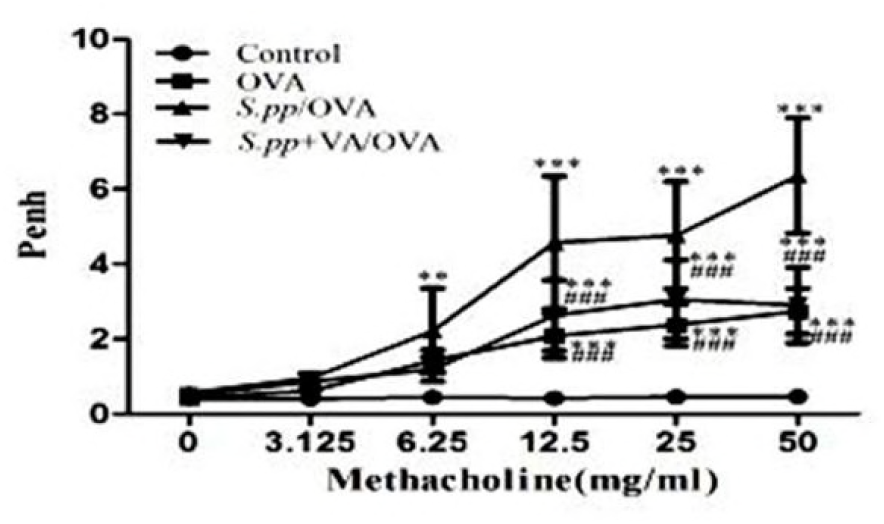
Vitamin A supplement after neonatal *S. pneumoniae* pneumonia alleviated AHR during AAD. Whole-body plethysmography in uninfected, non-allergic (Control), uninfected, allergic (OVA), neonatal infected, allergic (*S.pp* /OVA) and vitamin A supplement after neonatal infected, allergic (*S.pp* +VA/OVA) mice was conducted 24h following challenge with methacholine (n=6-8mice/group).**P<0.01, ***P<0.001 as compared with the control group,^###^P<0.001 as compared with *S.pp* /OVA group.

### Vitamin A supplement after neonatal *S. pneumoniae* pneumonia affect cytokines production during AAD

Twenty-four hours after the final challenge, BALF was obtained to detect the cytokines by ELISA. The productions of IL-4, IL-5, IL-13, IL-17A were significantly higher (P < 0.01), while IFN-γ was significantly lower in the *S.pp*/OVA group compared with the OVA group (P < 0.01).However, vitamin A supplement after neonatal *S. p*p dramatically decreased IL-4, IL-5, IL-13, IL-17A production and significantly increased IFN-γ production in the *S.pp*/OVA group. There was no significant difference of TGF-β production among OVA, *S.pp*/OVA and *S.pp*+VA/OVA groups (Table1).

**Table 1.**
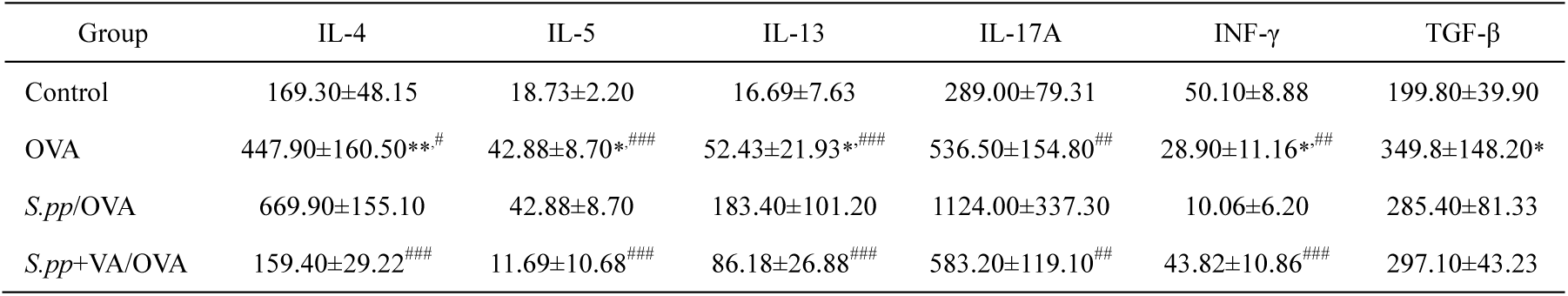
Cytokines productions in BALF during AAD (pg/ml) Concentrations of interleukin (IL)-4, IL-5, IL-13, IL-17A,interferon (IFN)-γ, and transforming growth factor (TGF)-β in the BALF of uninfected, non-allergic (Control), uninfected, allergic (OVA), neonatal infected, allergic (*S.pp* /OVA) and vitamin A supplement after neonatal infected, allergic (*S.pp*+VA/OVA) mice were measured by ELISA. Data are reported as mean ± standard error from three separate experiments (n = 6–8 mice/group). *P<0.05, **P<0.01, ***P<0.001 as compared with the control group,^#^P<0.05, ##P<0.01,^###^P<0.001 as compared with *S.pp* /OVA group.

### Vitamin A supplement after neonatal *S. pneumoniae* pneumonia alters the production of CD4^+^T cells during AAD

To determine the effects of vitamin A supplement after neonatal *S. p*p on the differentiation of CD4^+^T cells differentiation during AAD, flow cytometry was used to analyze the population of Foxp3^+^Treg, Th17, Th1and Th2 cells 24h after the final challenge. Data revealed that Foxp3^+^Treg and Th1 cells decreased significantly in *S.pp* /OVA group as compared with the OVA group (2.75±0.72% *vs* 5.30± 1.29%, P <0.001,1.13±0.33% *vs* 1.59±0.39%, P <0.05), while Th17 and Th2 cells increased (2.40±0.25% *vs*1.88±0.24%, P <0.001,1.50±0.17% *vs*1.05± 0.31%, P <0.05).

Vitamin A supplement after neonatal *S. pp* significantly increased Foxp3^+^Treg, Th1 cells production as compared with the *S.pp* /OVA group (5.64±2.11% *vs* 2.75± 0.72%, P <0.001, 2.27±0.36% *vs* 1.13±0.33%, P <0.01), while the number of Th17 and Th2 cells significantly decreased (1.93±0.12% *vs* 2.40±0.25%, P <0.001,0.78±0.31% *vs* 1.50± 0.17%, P <0.01). Thus, vitamin A supplement after neonatal *S.* pp promoted Foxp3^+^Treg and Th1 cells during AAD (Fig 6A-D).

**Fig 6.**
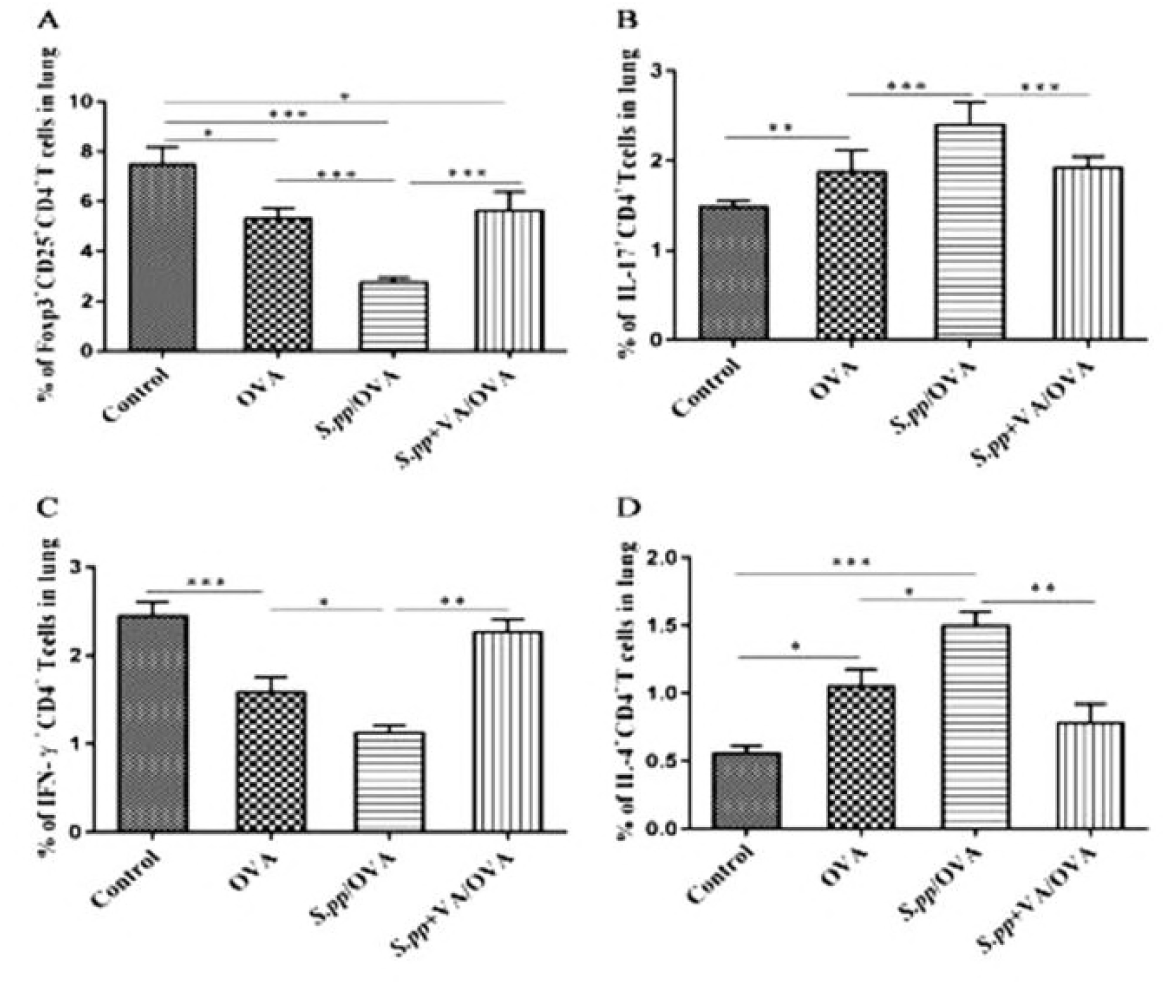
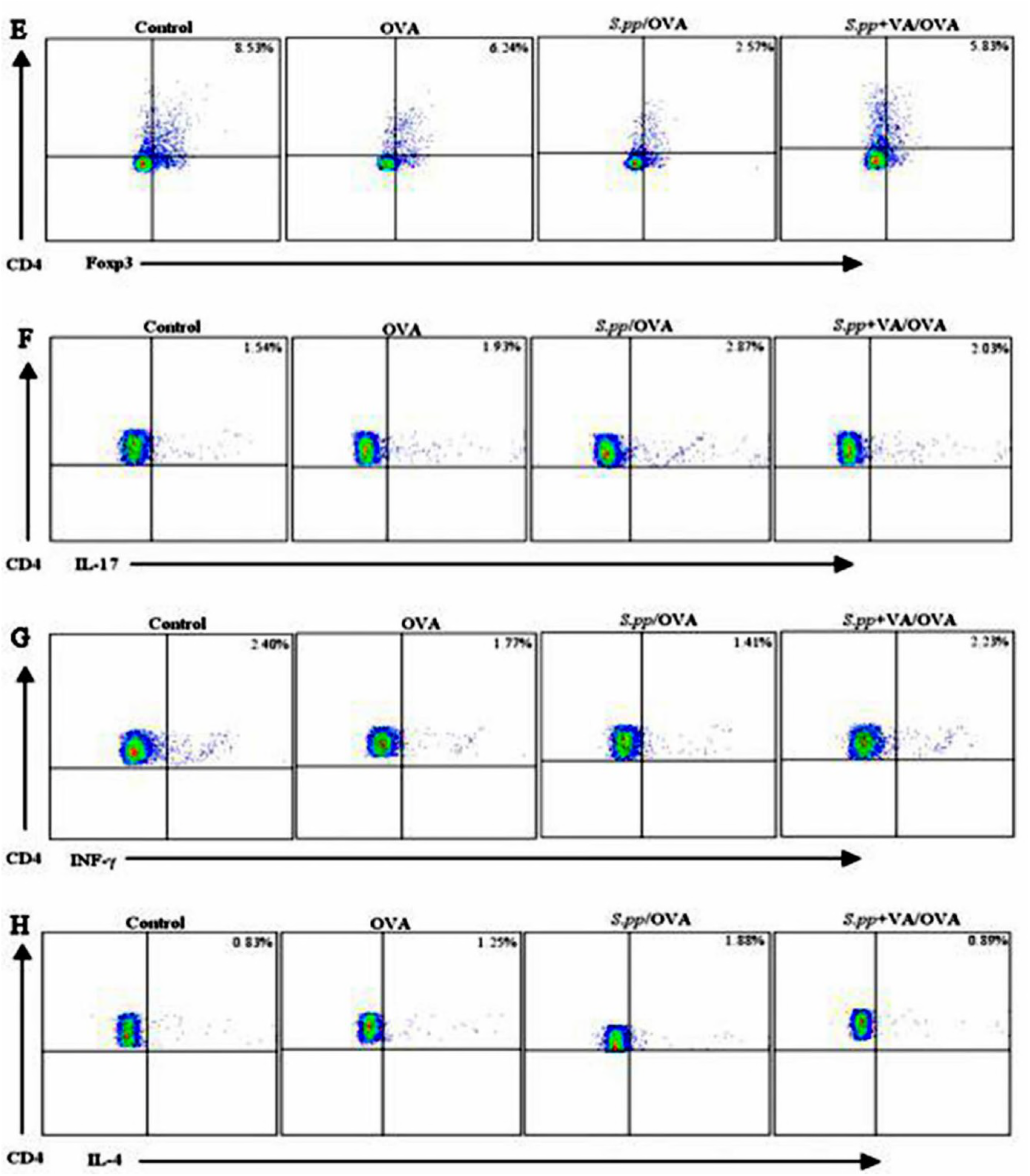
Vitamin A supplement after neonatal *S. pneumoniae* pneumonia altered CD4^+^T cells productions during AAD. Foxp3^+^Treg(A), Th17(B),Th1(C)and Th2(D) cells productions were measured in uninfected, non-allergic(Control), uninfected, allergic(OVA), neonatal infected, allergic(*S.pp* /OVA) and vitamin A supplement after neonatal infected, allergic (*S.pp*+VA/OVA) mice. The data (E–H), respectively, represent, percentages of positively stained cells of Foxp3+Treg, Th1, Th2 and Th17 within the lymphocyte gate of lung in BALB/c mice. (n=6–8 mice/group).*P<0.05, **P<0.01, ***P<0.001.

## Discussion

Asthma is one of the most common chronic diseases in children^[25]^. Epidemiological studies have demonstrated the association between early-life infections and subsequent asthma development^[26, 27]^. It is now well accepted that asthma is a heterogeneous syndrome with many clinical subtypes. Viral infections have been implicated in asthma pathogenesis as well as exacerbation^[28–31]^.

Infection by atypical pathogen (such as *Mycoplasma pneumoniae*) also appears to play important role in the induction and exacerbation of asthma both in children and adults^[32, 33]^. Recent studies suggest some bacterial infection have important role in asthma pathogenesis^[34, 35]^. Clinical study stated acute episodes of wheezing in some children are closely associated with bacterial infection^[32]^ and *S. pneumoniae* infection may increase the risk of asthma exacerbation^[36]^. Our previous study indicated that neonatal *S. pneumoniae* pneumonia promoted early adulthood allergic asthma development^[8]^. Pneumonia decreases vitamin A levels significantly in children under five years old^[13]^. In this study, we monitored vitamin A levels after *S. pp* and investigated the effect of vitamin A supplement post-infection on the development of allergic asthma. Our findings demonstrated that neonatal *S. pneumoniae* pneumonia induced lung vitamin A deficiency up to early adulthood, vitamin A supplement after neonatal *S. pneumoniae* pneumonia inhibited the recruitment of airway neutrophils and eosinophils, alleviated airway inflammation and decreased AHR during AAD. Vitamin A supplement not only promoted Foxp3^+^Treg and Th1 cells, but also inhibit Th2 cells production, which resulted in increased IFN-γ productions, decreased type? cytokines and IL-17A expressions during AAD. Our results indicate that vitamin A supplement after neonatal *S. pneumoniae* pneumonia alters lung CD4^+^T cell subsets and prevents subsequent allergic asthma development.

Others have reported that retinol was decreased in LPS or rhIL-6 treated infant rats^[18, 37]^, and in human infants and young children who have *S. pneumoniae* or other infections^[14, 38, 39]^. Similarly, we showed that neonatal *S. pneumoniae* pneumonia decreased lung vitamin A levels until early adulthood in mice. In contrast, Katherine et al^[9]^ found no significant differences in vitamin A level in serum and lung between *S. pneumoniae* infected and mock-infected adult mice, indicating that *S. pneumoniae* infection at different periods of life may induce different effects on vitamin A levels. Possible explainations for lung vitamin A deficiency after neonatal *S. pneumoniae* pneumonia include: 1) insufficient vitamin A storage in neonates; 2) increased vitamin A consumption: fast growth in neonates increases the need of vitamin A, repairing the damaged epithelial may increase vitamin A consumption^[13]^; 3) vitamin A deficiency may reduce retinol binding protein (RBP) production, leading to decreased mobilization of vitamin A from liver^[18]^; 4) ordinary food supply after neonatal pneumonia is insufficient to restore vitamin A concentrations^[40]^.

Epidemiologically vitamin A deficiency is common in asthmatic patients^[17, 41–43]^. Whether vitamin A deficiency induces asthma development or asthma cause vitamin A deficiency is not clearly understood. Growing evidence demonstrate that vitamin A directs immune cell differentiation and induces allergic disease. Intestinal studies in vivo and in vitro showed that sufficient retinoic acid (A kind of metabolites of vitamin A) can promote regulatory T cells productions^[44, 45]^. Akiko et al^[46]^ stated the differentiation of Foxp3^+^Treg from naïve CD4^+^T cell is decreased in vitamin A deficient mice. Animal studies suggest that sufficient vitamin A can suppress Th2 reaction and promote Foxp3^+^Treg and Th1 cells productions^[47–49]^. Consistent with these reports, our data showed that neonatal *S. pneumoniae* pneumonia reduced Foxp3^+^Treg and Th1 productions, increased Th2 cells during AAD, which aggravated allergic inflammation.

Vitamin A supplement after neonatal *S. pneumoniae* pneumonia may inhibit asthma development by inducing Foxp3^+^Treg and Th1 cells productions. Thus, our study indicates that neonatal *S. pneumoniae* pneumonia induces lung vitamin A deficiency alters local CD4^+^T cell differentiation during AAD and promots subsequent development of allergic asthma. Studies have reported negative correlation between vitamin A levels and the risk of asthma development^[16, 52, 53]^, while there has controversy between vitamin A supplement and asthma. Some studies demonstrated that sufficient vitamin A inhibit asthma or allergic disease by downregulating oxidative stress^[54]^, or via direct effects on the immune system^[49, 55–57]^. A recent study demonstrated dexamethasone therapy alone could not relieve allergic asthma airway epithelium injury, but combined with vitamin A promoted epithelium repair by down-regulating leucine zipper (GILZ) expression and activating MAPK-ERK signaling^[58]^. However, there are some studies also suggest association of vitamin A supplement with increased risk of asthma.

Some clinical studies found infants supplemented vitamins A or multivitamin showed increased risk of allergic disease^[59, 60]^. In addition, a study in Norwegian adults showed that daily intake of cod liver oil (rich in vitamin A) for ≥1 month significantly increased the incidence of adult-onset asthma^[61]^. One possible explanation for these inconsistencies is that excessive vitamin A may lead to its accumulation in the lung and hypervitaminosis A^[62]^. Hypervitaminosis A has been reported to be associated with airway hyperresponsiveness in mice model^[63]^. These findings suggest that hypervitaminosis A may increase the risk of asthma in response to allergens. Our study indicated that neonatal *S. pneumoniae* pneumonia resulted in lung vitamin A deficiency and promoted subsequent allergic asthma development. Vitamin A supplementation after neonatal *S. pneumoniae* pneumonia promoted Foxp3^+^Treg and Th1 productions, reduced Th2 cell expressions when exposed to the allergen, which resulted in AHR and alleviated infiltration by inflammatory cells infiltrationtion alleviation, and eventually inhibit adulthood allergic asthma development in mice model. Our finding may provide a novel strategy for the prevention of allergic asthma induced by *S. pneumoniae* pneumonia. While further researches are needed to explore the mechanisms in which neonatal *S.pneumoiae* pneumonia induces vitamin A deficiency. More studies are needed to clarify whether our results can to be extrapolated to other pathogens and other animals.

## Conclusions

Using a mouse model, we demonstrate that Vitamin A supplement after neonatal Streptococcus pneumoniae pneumonia alters the CD4^+^T cell subset and inhibits the development of early adulthood allergic asthma.

## Abbreviations

AAD: allergic airway disease
AHR: airway hyperresponsiveness
ANOVA: one-way analysis of variance
BALF: bronchoalveolar lavage fluid
HPLC: high performance liquid chromatograph
OVA: ovalbumin
PBS: phosphate buffered saline
*S. pneumoniae*: *Streptococcus pneumoniae*
*S.pp*: *S. pneumoniae* pneumonia

## Declarations

### Ethics approval and consent to participate

All experiments performed in mice were permitted by the Institutional Animal Care and Research Advisory Committee at the Chongqing Medical University. All experimental animals were used in accordance with the guidelines issued by the Chinese Council on Animal Care.

## Acknowledgments

We thank the Department of Laboratory Medicine, Key Laboratory of Diagnostic Medicine, Chongqing Medical University for providing the *S.pneumoniae* strain D39. We also thank the Experimental Animal Center at Chongqing Medical University for providing the BALB/c mice.

## Funding

This work was supported by the National Natural Science Foundation of China (81270086, 814700222), the Science and Technology Department of Chongqing (cstc2017jcyjB0160), the Science and Technology Department of Yuzhong district, Chongqing (20170120).

## Availability of data and materials

The datasets used and/or analyzed during the current study are available from the corresponding author on reasonable request.

## Authors’ contributions

Study design: YT, QT, YW, XP, ZL; conducting experiments: YT, QT, YW, XP; acquiring data: YC, QL, GZ, XT, LR; analyzing data: YT, QT, YW; writing the manuscript: YT, QT, XP. All authors read and approved the final manuscript.

## Consent for publication

Not applicable

## Competing interests

All authors: No reported con?icts. All authors have submitted the ICMJE Form for Disclosure of Potential Con?icts of Interest. Con?icts that the editors consider relevant to the content of the manuscript have been disclosed.

